# Cell-type and dynamic state govern genetic regulation of gene expression in heterogeneous differentiating cultures

**DOI:** 10.1101/2024.05.02.592174

**Authors:** Joshua M. Popp, Katherine Rhodes, Radhika Jangi, Mingyuan Li, Kenneth Barr, Karl Tayeb, Alexis Battle, Yoav Gilad

## Abstract

Identifying the molecular effects of human genetic variation across cellular contexts is crucial for understanding the mechanisms underlying disease-associated loci, yet many cell-types and developmental stages remain underexplored. Here we harnessed the potential of heterogeneous differentiating cultures (**HDCs**), an *in vitro* system in which pluripotent cells asynchronously differentiate into a broad spectrum of cell-types. We generated HDCs for 53 human donors and collected single-cell RNA-sequencing data from over 900,000 cells. We identified expression quantitative trait loci in 29 cell-types and characterized regulatory dynamics across diverse differentiation trajectories. This revealed novel regulatory variants for genes involved in key developmental and disease-related processes while replicating known effects from primary tissues, and dynamic regulatory effects associated with a range of complex traits.

## Introduction

Decoding the molecular consequences of genetic variation is a central goal in human genetics. With the advent of genome-wide association studies, a vast array of genetic variants associated with diseases have been uncovered. These predominantly lie in non-coding regions of the genome, suggesting primarily regulatory mechanisms ^1^. This insight has spurred a surge in mapping expression quantitative trait loci (eQTLs) to understand how the disease-associated genetic variants influence gene expression levels. Despite significant strides made by several large-scale projects such as the GTEx Consortium to map eQTLs ^2–7^ , a comprehensive understanding of the molecular impacts of disease-associated loci remains elusive, in part due to the context-dependent and dynamic nature of gene regulation ^8–11^.

Gene regulation varies by contexts including cell-type, temporal stage, and environment. This poses a formidable challenge for human studies that seek to characterize the gene regulatory basis for complex traits ^12–14^. Studies using postmortem human tissues, while delivering important insight, often fall short of capturing the full spectrum of dynamic regulatory effects because they reflect predefined adult tissue contexts, and because most studies have utilized bulk sequencing. Recent advances in single-cell technologies have started to shift this paradigm by enabling researchers to collect a heterogeneous biological sample and disentangle context-specific regulatory variation through downstream analysis of single-cell molecular phenotypes ^15–20^. Still, many contexts are difficult to sample from healthy human donors, particularly dynamic contexts where we would like to capture multiple time points from the same individual. This would be nearly impossible for an inaccessible tissue. This has motivated the use of differentiation protocols for *in vitro* cell cultures, which have offered access to dynamic regulatory effects, including fleeting effects present only at intermediate stages of differentiation^21^. These *in vitro* systems are each imperfect reflections of human cell biology, but the activation of a range of relevant *cis*-regulatory elements can reveal the effects of variants within them even without completely recapitulating *in vivo* cellular state. Indeed, studies in these systems have captured regulatory effects of numerous disease-associated loci and variants near genes involved in developmental processes ^15,16,21^. However, most protocols would require separate experimental setups for each cell-type or perturbation of interest, making it difficult to efficiently explore the space of disease-relevant contexts.

In this study, we explored gene regulation across diverse cellular contexts using heterogeneous differentiating cultures. We introduce the terminology “heterogeneous differentiating cultures (HDCs)” as a broad descriptor encompassing a class of related *in vitro* models that can be used to explore diverse cellular contexts efficiently. Here, we focus on unguided HDCs, which are based on embryoid body systems using an extended culturing time to consistently generate dozens of cell-types derived from all three developmental germ layers ^22,23^. We have also developed guided HDCs, which push iPSCs toward certain lineages to enrich for multiple cell types within a particular tissue and offer the ability to pursue more targeted questions. While both unguided and guided HDCs differ from *in vivo* cellular biology as expected, we have demonstrated in previous studies and here that the expression profiles and genetics effects found in HDCs overlap those found using primary cell types and tissues ^24,25^.

Here, we generated unguided HDCs from a panel of 53 human iPS cell lines and measured gene expression at single-cell resolution in over 900,000 cells. We mapped eQTLs in 29 cell-types, including many that have never before been characterized at the population level in humans, and identified dynamic genetic effects on gene regulation that vary with respect to diverse differentiation trajectories and gene programs.

## Results

### Heterogeneous differentiating cultures generate diverse cell-types

We established unguided HDCs from the iPSCs of 53 unrelated Yoruba individuals from Ibadan, Nigeria (YRI) (Fig. 1A; Methods). Briefly, we formed HDCs in batches of 4-8 individuals and maintained them in culture for 21 days (Supplementary Table S1). Within each batch, we formed, maintained, and dissociated HDCs in parallel. After dissociation, we multiplexed samples from each individual in equal proportions in preparation for single-cell RNA sequencing, targeting a depth of 100,000 reads per cell. After quality control, filtering, and de-multiplexing, we retained data from 909,536 high-quality cells (median 1,241 cells per individual per replicate; median 21,990 UMI counts per cell).

**Fig. 1:**
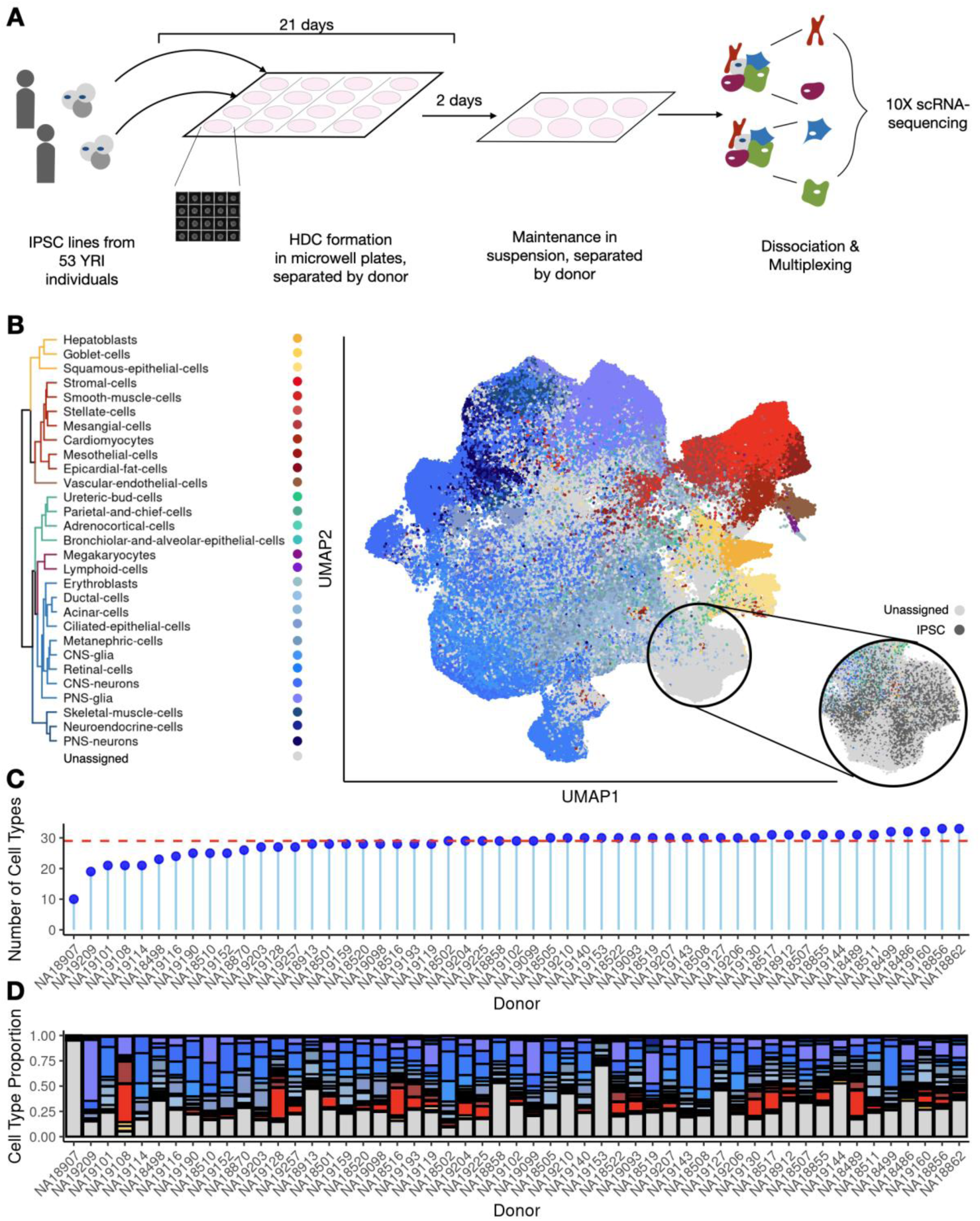
Heterogeneous differentiating culture (HDC) panel from 53 human iPSC lines. (A) Schematic illustration of the data generation process. (B) UMAP embedding of over 900,000 HDC cells, annotated using the fetal cell atlas. Inset shows annotation of a primarily unassigned group of cells with an augmented classifier trained with pluripotent cells. (C) Most cell lines generate most cell-types with sufficient coverage for inclusion in QTL analyses. (D) Cell-type proportions vary between lines.

To initially assess cellular diversity in HDCs, we developed a cell-type classifier based on data and annotations from the fetal cell atlas ^26^. We used a curated set of 33 high confidence cell-type labels (Methods, Fig. S1) that span the three main germ layers (Fig. 1B). We assigned 651,129 cells (72% of all cells) to one of these cell-types based on gene expression signatures (Supplementary Table S2). Using this approach, 28% of the cells remained unclassified. Of the 33 cell-types, 29 are represented with a minimum of 5 cells in at least 25 individuals. While the proportion of cells of each type varied between individuals (Fig. 1C), 52 of 53 individuals have data from least 5 cells from most cell-types (median 29 of 33 cell-types per donor, Fig. 1D).

Some of the unannotated cells express markers of pluripotency, suggesting that asynchronous differentiation within HDCs enabled us to collect pluripotent cells alongside partially and fully differentiated cell-types. Indeed, manually adding a marker gene set for iPSCs ^27^ to the list of cell-type signatures enabled us to classify 21,370 previously unannotated cells as iPSCs (Fig. 1B). Since this iPSC signature was obtained from a separate reference, and since eQTLs in iPSCs have previously been thoroughly characterized ^28,29^, we focused on the 29 common fetal cell atlas cell-types in the subsequent cell-type-stratified eQTL analysis, filtering to the 52 donors that successfully differentiated into diverse cell-types (Supplementary Table S3). We re-incorporate these pluripotent and unannotated cells in later analyses that focused on evaluating regulatory dynamics across the HDC system.

### eQTLs across cell-types

We mapped genetic effects on gene regulation in each of the 29 discretely defined fetal cell atlas cell-types. To mitigate the effects of noise inherent to single-cell data, we aggregated single-cell expression into pseudobulk such that each observation represents all individual cells from a single donor/cell-type combination ^30,31^ (Methods). This aggregation step also allowed us to take advantage of well-established methods for eQTL mapping using bulk RNA-sequencing data.

We initially performed *cis* eQTL mapping specifically in each cell-type, limiting our analysis to the 29 annotated cell-types with at least 5 cells from at least 25 donors (Fig. 2A). We included expression principal components for data from each cell-type as covariates, to control for hidden factors driving global expression variability, including batch effects ^32^. Across all cell-types, we identified a total of 31,179 eQTLs (associated with 2,114 eGenes) at a global q-value cutoff of 0.05 ^33^. 79% of these HDC eQTLs were previously identified by GTEx (Fig. 2B); that is, 6,572 of our eQTLs have not been identified before. The HDCs include many developing cell-types not found in GTEx adult tissues. Indeed, the subset of eGenes regulated by novel eQTLs, identified in HDCs but not in GTEx, were enriched for several developmental processes, including tissue development (OR=2.08, two-sided Fisher’s exact test p=4.5e-5), central nervous system development (OR=2.31, p=1.2e-4), and circulatory system development (OR=2.32, p=1.3e-4) (Supplementary Table S4). The clustering of novel eQTLs upstream of transcription start sites (Fig. S2) indicates that these enrichments are not an artifact due to false positive associations near developmental genes that are relatively depleted for eQTLs in GTEx ^11^.

**Fig. 2:**
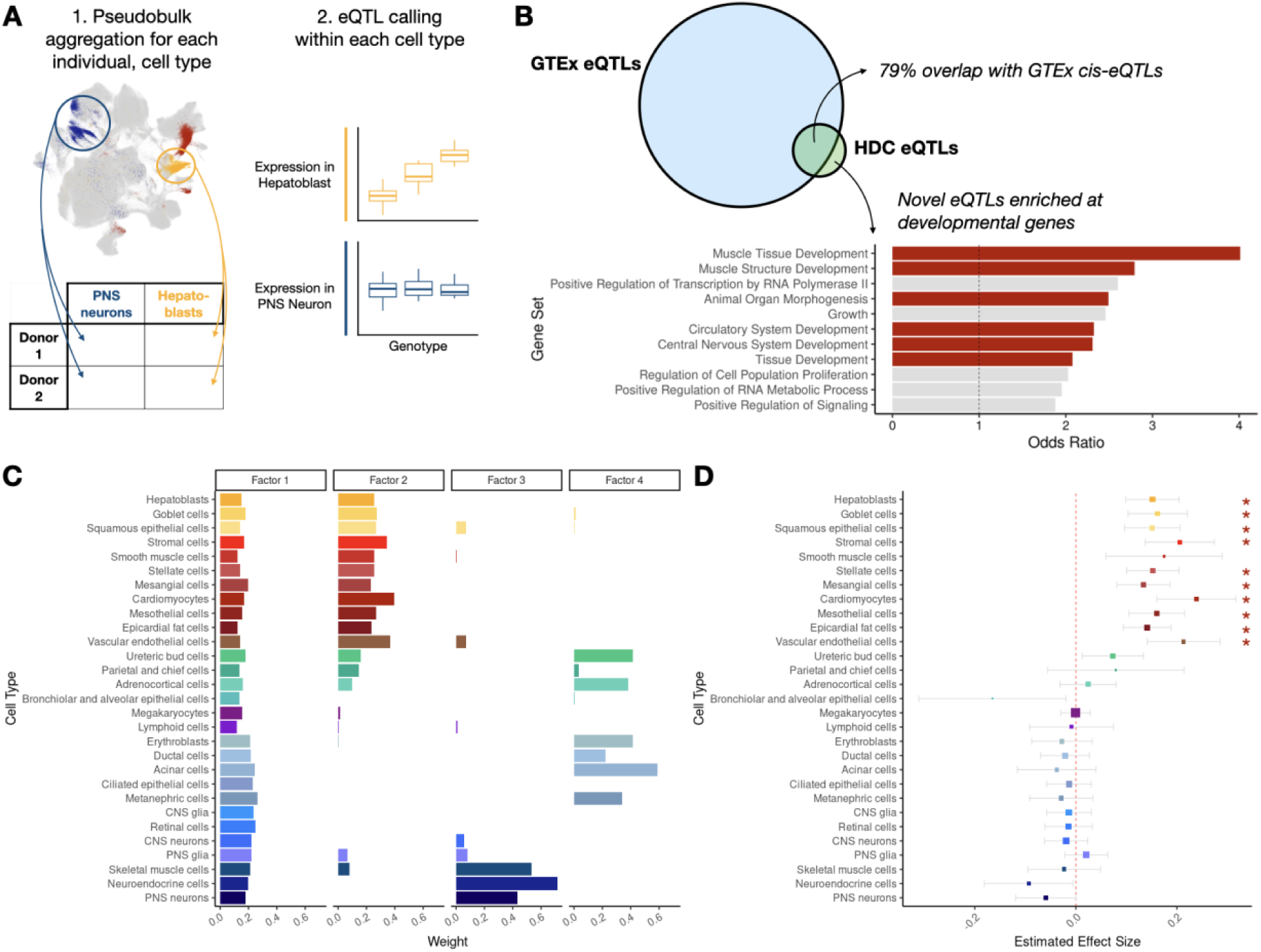
eQTL calling across 29 cell-types. (A) To perform eQTL calling, UMI counts from cells from the same donor and cell-type were aggregated into a pseudobulk sample, and eQTL calling was performed separately in each cell-type. (B) Comparison of HDC eQTLs to GTEx eQTLs. Bar plot shows odds ratio of 11 GO biological processes significantly enriched for HDC eGenes with no eQTLs overlapping GTEx hits (FDR <= 0.05, background gene set all HDC eGenes). Developmental processes are highlighted in red. (C) Patterns of regulatory effects learned through matrix decomposition of eQTL effect sizes in each cell-type. Beyond the densely loaded factor 1, remaining factors partition cell-types of similar developmental origin. (D) Metaplot of eQTL for the gene *SH3PXD2B* at rs10042482. Boxes are centered at the estimated posterior mean effect in the given cell-type, error bars show +/- posterior standard deviation, box size indicates precision (1/squared posterior standard deviation). This eQTL was not detected in GTEx, or the univariate (cell-type-by-cell-type) analysis. (*) indicates significance at local false sign rate of 0.05.

Within each cell-type, we identified a median of 2,099 eQTLs (maximum of 15,064 eQTLs in PNS glia cells) in a median of 126 eGenes (maximum of 919 eGenes in PNS glia cells) (Supplementary Table S5). The number of eQTLs detected in each cell-type is correlated with the median number of individual cells from which we have data per cell-type, across individuals (Fig. S3), suggesting that power to detect eQTLs is limited by both sample size and the number of captured individual cells. To account for incomplete power to detect eQTLs in any given cell-types, and to assess the extent of eQTL sharing between cell-types, we analyzed the data using multivariate adaptive shrinkage (mash) ^34^. By borrowing information across cell-types and genes, mash allows us to detect weak signals that emerge repeatedly in different cell-types with greater confidence, thereby improving power. More generally, mash can be used to identify major patterns of heterogeneity and sharing between cell-types.

The four strongest regulatory patterns revealed when we used this approach depict broadly shared regulation among all cell-types, and then among subsets of developmentally related cells. Of these more specific patterns, the first corresponds to endoderm- and mesoderm-derived cell-types, the second to ectoderm-derived cell-types, and the third to the cluster of epithelial cell-types which may correspond to neuroepithelium, which would not be expected to appear in our fetal cell atlas reference (Fig. 2C). We incorporated these candidate regulatory patterns alongside several ’canonical’ patterns such as single cell-type effects into a prior distribution that guides hypothesis testing (Methods).

Using mash, we detected an additional 56,614 eQTLs corresponding to 4,973 eGenes (Supplementary Table S6). The example of *SH3PXD2B* (Fig. 2D), a gene thought to be important for cardiac and skeletal development ^35^, highlights the utility of considering structured regulatory patterns in the analysis of heterogeneous single-cell data. Indeed, no significant eQTLs for *SH3PXDB2* were found in GTEx, nor the univariate analysis considering one HDC cell-type at a time (Fig. S4). Specifically, when we leveraged regulatory patterns shared by cell-types of similar developmental origin, we found a significant *cis* eQTL for *SH3PXDB2* (local false sign rate; lfsr <= 0.05) in nearly all endoderm- and mesoderm-derived cell-types.

### Dynamic genetic regulation along diverse differentiation trajectories

Next, we took a different approach to characterize how interactions between genetic variants and the cellular environment impact gene expression levels. To do so, we abandon discretely defined cell-type labels in favor of more continuous and nuanced representations of cellular variation.

We were particularly interested in characterizing temporally dynamic genetic effects, including effects that fluctuate during cellular development. As cells within HDCs differentiate asynchronously, our current dataset captures continuous gene regulatory variation along multiple developmental trajectories. To validate our approach for defining differentiation trajectories in HDCs, we began by focusing on the cardiomyocyte lineage, because we had previously collected single-cell data from directly differentiated cardiomyocytes using a time course study design, which can be used as the ground truth for both gene expression and regulatory dynamics. First, we compiled manually curated gene lists containing marker genes from each stage of cardiomyocyte differentiation (Methods, Supplementary Table S7) ^17^. We used a cell-scoring tool (scDRS) ^36^ to identify subpopulations of cells with enriched expression for gene sets specific to the cardiomyocyte trajectory, and applied principal components analysis to infer pseudotime (Methods, Fig. 3A) ^16,37^. The first expression principal component using data from these cardiomyocyte trajectory cells offers a reasonable pseudotime metric (Fig. 3B), as it captures sequential expression of marker genes for each stage of cardiomyocyte differentiation (*NANOG*, an IPSC marker gene; *MIXL1*, mesendoderm; *MESP1*, mesoderm; *GATA4*, cardiac progenitor; and *TNNT2*, cardiomyocyte) ^38^. Following our previous approach ^21^, we identified 709 linear dynamic eQTLs (in 47 eGenes) along the cardiomyocyte trajectory at a genome-wide FDR 0.1, using EigenMT to control for multiple testing burden within each gene (Methods). The majority of these effects were replicated in the directly differentiated cardiomyocytes, where the sample size was far smaller (n=19; *π*_1_ replication rate = 0.58) ^17^.

**Fig. 3:**
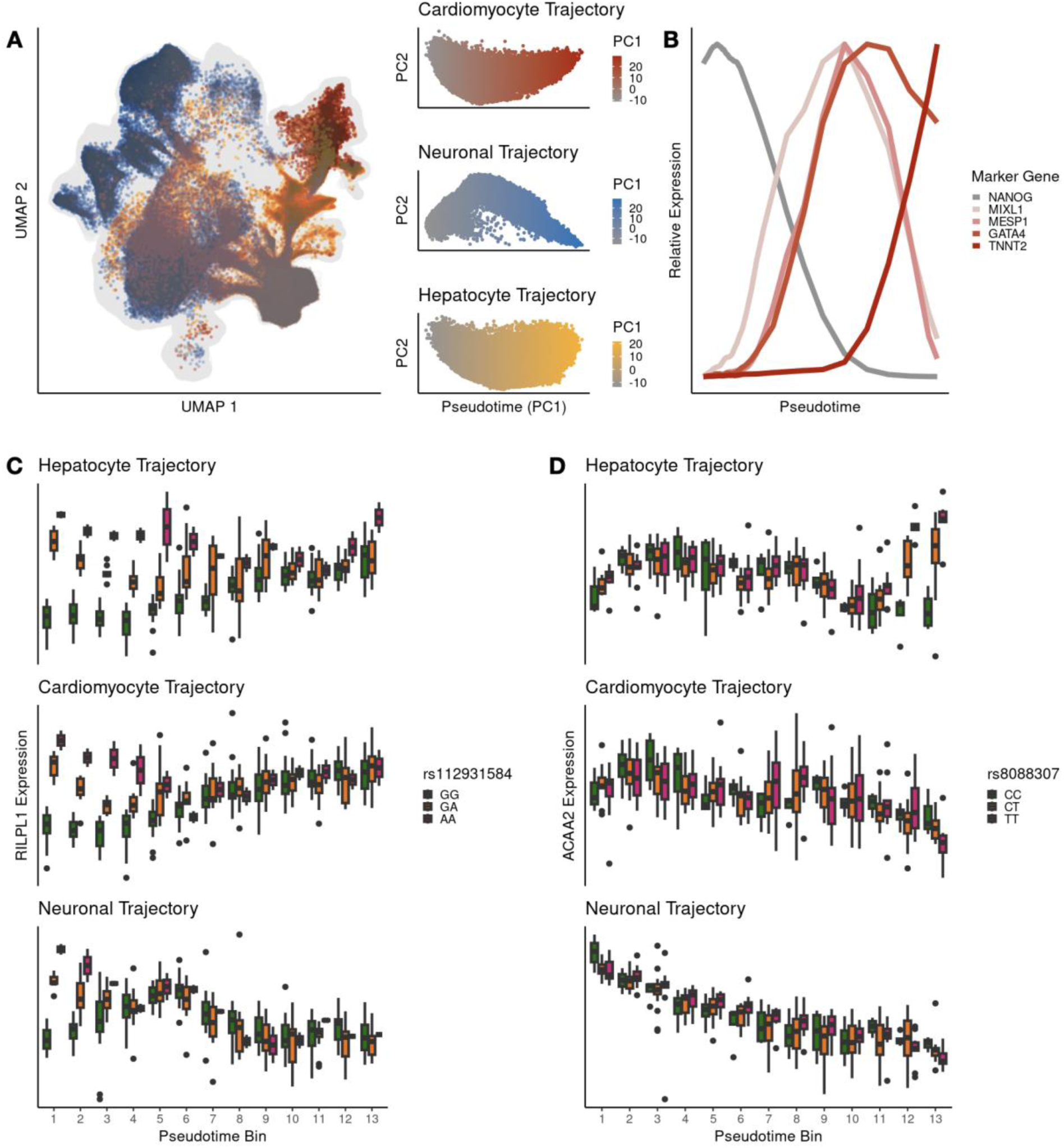
Dynamic eQTL calling along all three germ layers. (A) Trajectory isolation extracts cells mapped to three differentiation trajectories from the full dataset (left). The first expression principal component in each trajectory is used as a measure of pseudotime (right). (B) Relative (min-max normalized) expression of 5 key marker genes displaying sequential expression with respect to pseudotime, (C) Early dynamic eQTL for the gene *RILPL1*, shared across all three differentiation trajectories (center line, median normalized expression; box limits, upper and lower quartiles; whiskers, 1.5x interquartile range; points, outliers). (D) Late dynamic eQTL for the gene *ACAA2*, specific to the hepatocyte differentiation trajectory (top).

After validating our approach in the cardiomyocyte trajectory, which is derived from mesoderm, we examined neuronal and hepatic differentiation trajectories, which respectively represent the ectodermal and endodermal germ layers (Supplementary Table S8-S9). At a genome-wide FDR of 0.1, we found 1,965 dynamic eQTLs (166 eGenes) along the neuronal differentiation trajectory and 472 dynamic eQTLs (44 eGenes) along the hepatocyte differentiation trajectory (Supplementary Table S10).

We further categorized the dynamic eQTLs we identified in these three developmental trajectories into early (40%), late (57%), and switch (3%) effects, excluding three genes where eQTL classification diverged between trajectories (see Methods). For example, *RILPL1*, which is thought to regulate cell shape and polarity ^39^, is associated with an early dynamic eQTL in all three trajectories (Fig. 3C), while *ACAA2*, which encodes a protein involved in fatty acid metabolism in the liver, is associated with a late eQTL only in hepatocytes (Fig. 3D). While 77% of late and switch dynamic eQTLs overlap previously discovered effects listed in the GTEx catalog, similar to the proportion of overlapping cell-type eQTLs, the overlap with GTEx drops to 60% for early dynamic eQTLs. That is, eQTLs identified only in the early developing cell-types of each trajectory are markedly less likely to be observed in GTEx adult tissue eQTLs.

### Resolving complex regulatory interactions using topic analysis

Cell-type and differentiation stage represent the most salient aspects of cellular identity in our dataset. Regulatory changes along differentiation and between cell-types lead to strong gene expression differences, such that straightforward application of unsupervised machine learning methods (such as clustering and principal components analysis) to gene expression will first stratify cells based on these features. However, as they differentiate, cells are simultaneously engaging in a wide array of dynamic processes, including growth, division, and signaling. Many of these processes have the potential to alter gene regulation across different trajectories, and perhaps also orthogonally to the effects of cell-type or differentiation stage.

To obtain a richer description of cellular identity, and potentially resolve contrasts in cellular context beyond the discrete definition of cell-type and differentiation stage, we performed topic modeling of HDC expression data. In this framework, transcriptional variation is decomposed into a fixed number of ‘cellular topics’ represented by functional gene modules. We used FastTopics to identify 10 topics in the HDC dataset, aggregating cells into pseudocells (small clusters of transcriptionally similar cells) to mitigate noise and improve computational efficiency (Methods) ^40^.

Several topics appeared to overlap with cell-type and/or germ layer labels (Fig. 4A), essentially recapitulating the results of cell-type annotation: topic 1 was heavily loaded across endoderm-derived cell-types, topic 2 across mesoderm-derived cell-types, topic 5 on glial cells and topic 6 on ectoderm-derived cell-types. Two more topics encode developmental stage-specific information: topic 4 is most highly loaded on pluripotent cells, and topic 7 is most heavily loaded on cells at intermediate stages of neuronal differentiation (Fig. S5). This reinforces that cell-type identity and developmental stage are the primary drivers of transcriptional variation, as mentioned above. Other topics, however, appeared to stratify cells based on gene programs that are less dependent on cell-type or trajectory. For example, gene set enrichment analysis suggests that topic 8 appears to track the signature of cell cycle in our data (Supplementary Table S11). We confirmed this by directly estimating cell cycle phase for all pseudocells (Fig. 4B) ^41^. Topic 10 corresponds to a ciliary gene program that is shared across many cell-types represented in this dataset.

**Fig. 4:**
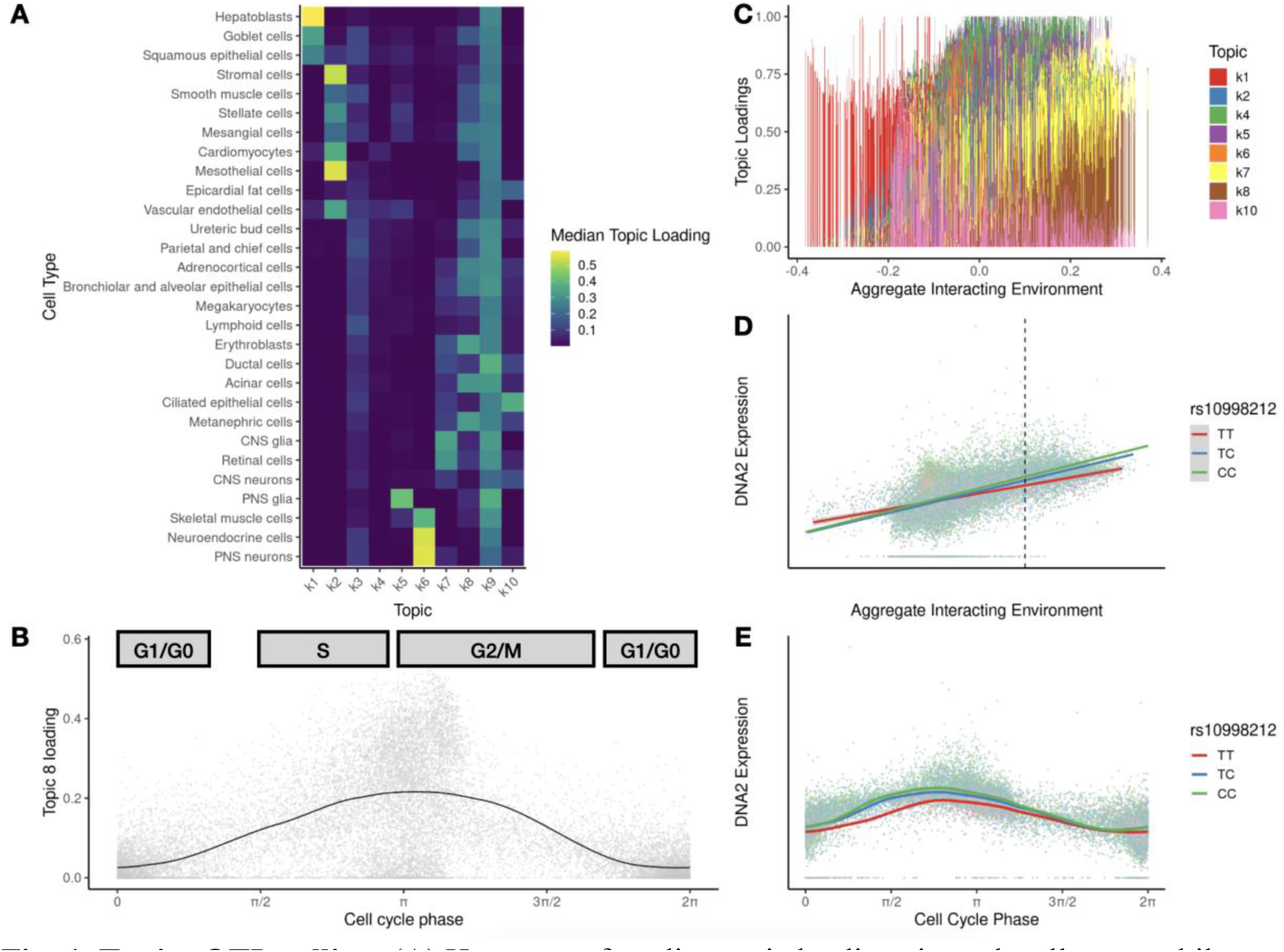
Topic eQTL calling. (A) Heat map of median topic loadings in each cell-type: while some topics are specific to one or a few cell-types, others are associated with processes broadly shared across cell-types. (B) Topic 8 is associated with cell cycle phase, with the highest loadings on cells in G2/M stage (phase ). (C-D) *DNA2* has an eQTL whose effect varies over the course of the cell cycle. CellRegMap identifies this effect by testing for interactions between genotype and any linear combination of latent topics (D); aggregate interacting environment is the linear combination which maximizes this interaction. (E) Visualizing the effect with respect to directly estimated cell cycle phase offers a more interpretable view of the regulatory dynamics, with maximal effect occurring in S phase.

We next sought to identify genetic variants with regulatory effects that are specific to the expanded set of cellular processes that were captured as topics (i.e., topic eQTLs). We limited this analysis to the 8 topics described above, as these were the most interpretable (Methods) ^42^. We used CellRegMap to map topic eQTLs ^43^. Instead of assessing each topic in isolation, CellRegMap jointly considers all linear combinations of topics, enabling us to simultaneously test for a wide range of genetic interactions with the cellular environment.

Since this powerful testing scheme is computationally expensive, we limited the topic interaction testing to the 77,550 eQTLs identified in the mash analysis. We identified a total of 157 genes with a topic eQTL (Supplementary Table S12). For example, *DNA2* has a topic eQTL whose effect is correlated with the apparent cell-cycle topic (topic 8; Fig. 4C-E). *DNA2* encodes a helicase protein involved in maintaining mitochondrial and nuclear DNA stability during DNA replication and repair. Visualizing this eQTL with the more clearly defined cell cycle phase inferred by tricycle ^41^ shows that this regulatory effect is largest during S phase, when DNA replication occurs. Another compelling example is the topic eQTL associated with *AXDND1*, with an effect that is correlated with the ciliary topic (topic 10; Fig. S6). *AXDND1* is thought to encode a component of axonemal dyneins, which drive ciliary beating to enable cell motility and extracellular fluid flow ^39,44^. This analysis provides granular functional insight that is obscured by the standard catalog-based eQTL mapping approach.

### HDC eQTLs help reveal the functional context of GWAS loci

Our eQTL mapping approach leverages the extensive cellular heterogeneity present in HDCs to identify gene regulatory variants whose effects are most prominent in underexplored contexts, ranging from early developmental time points to gene programs that are active across multiple cell states. Context-specific genetic effects could potentially shed light on disease-associated loci that have thus far failed to display regulatory potential in steady-state adult tissue datasets. As a preliminary appraisal of our granular, multi-dimensional framework for characterizing gene regulation in HDCs, we investigated the intersection between the context-dependent effects we identified and disease-associated loci.

We first turned to schizophrenia, a highly polygenic disease with known contributions in early development ^45^. We found that eQTLs identified in the cell-type-by-cell-type analysis displayed stronger association with disease risk than a random set of variants (Fig. 5A). This inflation is driven in part by previously uncharacterized eQTLs: 42 of out 221 HDC eQTLs (corresponding to 8 out of 16 eGenes) displayed genome-wide significant association with schizophrenia (P <= 5e-8), but do not overlap with or tag (at LD R^2^ >= 0.5) any significant eQTL identified by GTEx. These disease-associated novel eQTLs include one particularly compelling example identified in PNS Glia cells for the gene *BCL11B*, a transcription factor involved in neurodevelopment (Fig.5B) ^46–49^.

**Fig. 5:**
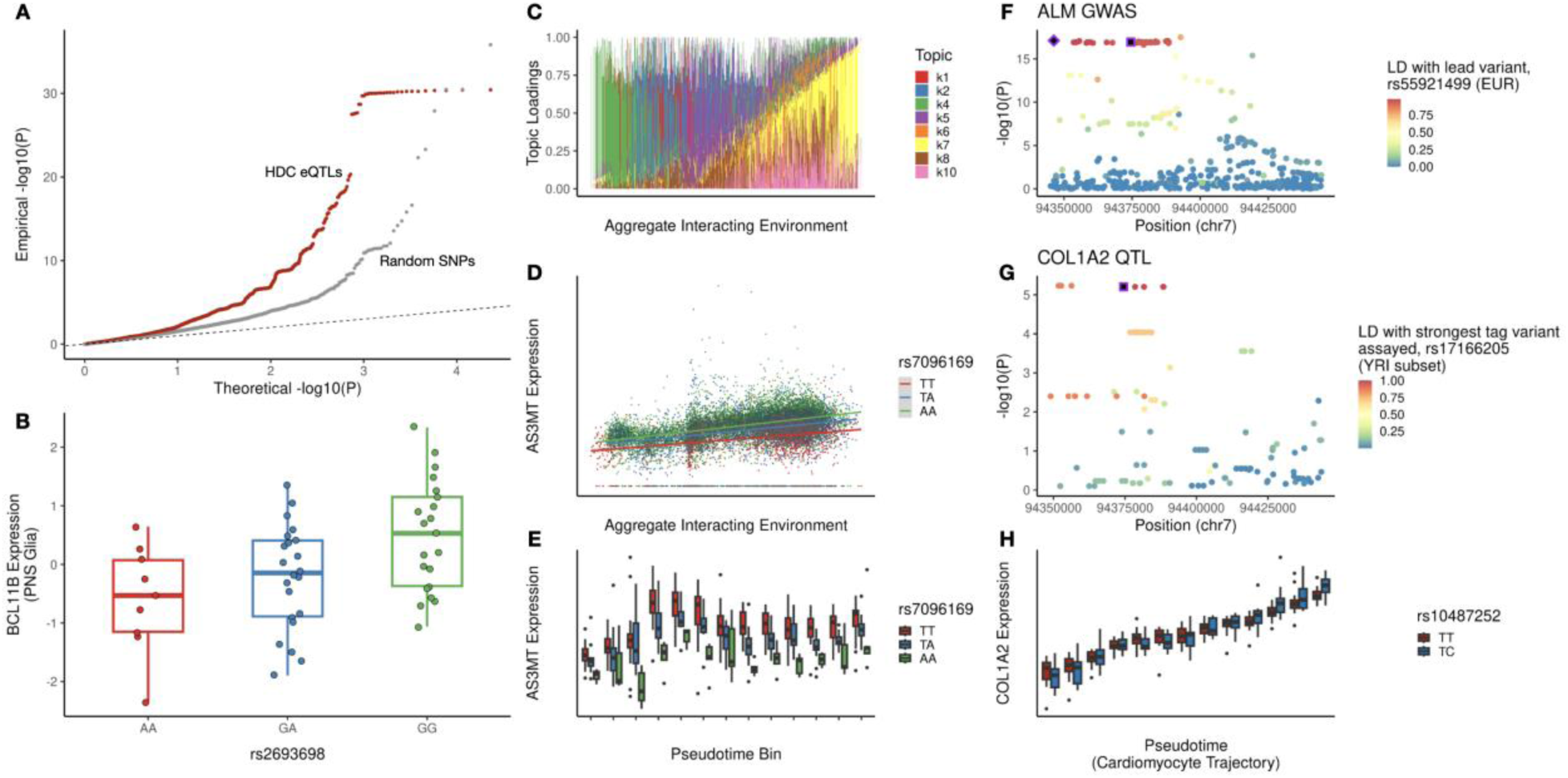
Exploring regulatory impacts of GWAS loci with HDCs. (A) HDC eQTLs (red) display inflation of small p-values from the GWAS for schizophrenia, compared to random SNPs (gray). (B) This inflation is driven in part by novel regulatory effects discovered in this system, such as the example shown: an eQTL for the gene *BCL11B* discovered in PNS Glia (center line, median normalized expression; box limits, upper and lower quartiles; whiskers, 1.5x interquartile range; points, individual observations). (C) A topic-interaction eQTL at schizophrenia-associated locus rs7096169, a topic eQTL for *AS3MT*, draws the greatest contrast between the two endpoints of neuronal differentiation (topic 4, green, highest in pluripotent cells, and topic 6, purple, highest in neuronal cells), and the intermediate stages (topic 7, yellow). (D) CellRegMap detects a significant interaction between genotype and this aggregate interacting environment on expression levels of the gene *AS3MT*. (E) Visualizing this effect with respect to the more intuitive pseudotime measurement of neuronal differentiation state clarifies the nonlinear dynamic effect of cell state on the eQTL. (F) Dynamic eQTL over the course of cardiomyocyte differentiation at an appendicular lean mass (ALM)-associated genetic variant. (G) Locus plot displaying significance of association with appendicular lean mass, points are colored by LD with the lead variant, shown with black diamond. As this variant has low frequency in the YRI panel used in the present study and was not tested for QTL effects, the assayed variant in strongest LD with the lead is additionally highlighted in the black square. (H) Locus plot displaying significance of the interaction between genotype and pseudotime along the cardiomyocyte trajectory on *COL1A2* expression.

Beyond mapping of individual genetic variants to candidate target genes, a more thorough characterization of regulatory dynamics offers insight into when and where the effects of a genetic variant may be most important. For example, rs7096169 is a known eQTL for *AS3MT* in several tissues, as well as a genome-wide significant schizophrenia risk locus. In HDCs, we found that this eQTL displayed a topic interaction effect, with the largest effect found in a topic associated with intermediate stages of neuronal development (topic 7), rather than either of the topics associated with an endpoint of neuronal development (topics 4 and 6, Fig. 5C-D). We can clearly observe this variant’s nonlinear dynamic regulatory effect when viewed with respect to pseudotime along the neuronal trajectory (Fig. 5E). While this triangulation of variant, gene, and context has previously been reported ^50,51^, such efforts have historically required scanning large-scale databases of regulatory effects in adult or immune contexts, then establishing an *in vitro* platform to evaluate expression during differentiation into a single cell-type selected *a priori*. The unification of an expansive set of cellular contexts at various stages of differentiation has the potential to accelerate this process, particularly when it is feasible to explore population-level genetic differences.

With this in mind, we next focused on novel interaction eQTLs (i.e., temporally dynamic eQTLs and topic eQTLs that are not found in GTEx, and did not tag a GTEx eQTL in any tissue at R^2^ >= 0.5). We used the Open Targets Genetics database to search for intersection between these novel regulatory effects and GWAS loci ^52^. We found 62 genes with novel interaction eQTLs (ieGenes) displaying genome-wide significant association (P <= 5e-8) with at least one disease-associated locus: 26 neuronal dynamic ieGenes, 8 hepatocyte dynamic ieGenes, 10 cardiomyocyte dynamic ieGenes, and 24 topic ieGenes (Supplementary Table S13). We highlight an example of a previously unknown dynamic eQTL for the collagen gene *COL1A2*, where the effect switches direction over the course of cardiomyocyte differentiation, suggesting a regulatory element with contrasting effects over time, or the convergence of multiple dynamic effects in LD with the variant shown (Fig. 5F-H). This eQTL tags a variant that is associated with appendicular lean mass, a measure of the musculature in arms and legs.

## Discussion

The exploration of unguided heterogeneous differentiating cultures (HDCs) in this study reveals an array of context-specific eQTLs that remained undetected in datasets such as GTEx, despite its comprehensive analysis of dozens of postmortem tissue types from hundreds of adult individuals. Our observations underscore the nuanced complexity of genetic regulation, which operates within highly specific cell-types, states, and temporal contexts. By exploring cell-types, trajectories, and programs that are difficult to sample *in vivo* and have never been studied from a population-level sample in humans, our work provides further support for the critical importance of context in understanding the regulatory mechanisms influencing disease.

We have demonstrated here that the HDC system offers valuable insights into gene regulation despite a lack of spatial organization mirroring in vivo tissues. Most of the cell-type eQTLs discovered here overlap loci with regulatory function previously reported in primary tissue samples from the GTEx project. The HDC eQTLs that do not display this overlap demonstrate the expected clustering around the transcription start sites of genes, but are enriched at known ’blind spots’ in existing resources: namely, the genes involved in a wide variety of developmental processes. Prioritizing flexibility enables us to activate a broader range of *cis-*regulatory elements which may be dormant in more accessible contexts.

Among the noteworthy findings from our HDC exploration is the identification of regulatory roles for dozens of disease-associated mutations that had not been linked to gene regulation in existing human eQTL studies. This highlights an important advantage of utilizing HDCs, which is the capacity to investigate a wide range of cellular contexts, some of which are rare or otherwise inaccessible in typical human samples. Enhanced by the detailed resolution of single-cell data and refined through trajectory inference and topic modeling, our approach enables the examination of not only distinct cell-types and states but also subtler functional contexts that drive changes in gene regulation, and the corresponding context-specific eQTLs.

The application of topic modeling to single-cell RNA-seq from HDCs has allowed us to traverse beyond traditional analyses confined by cell-type or overall gene expression correlations. Topic modeling has revealed hidden layers of regulatory variation driven by dynamic processes such as cell division and ciliary activity that occur across multiple cell types and trajectories. By uncovering these additional dimensions of cellular identity, topic modeling has proven essential in identifying specific functional contexts that remained cryptic within the more conventional frameworks of single-cell classification.

To bridge the gaps in our understanding of the genetics of gene regulation, it is imperative to move beyond the static snapshots provided by adult tissues that have dominated eQTL studies in humans. Heterogeneous differentiating cultures facilitate this expansion by offering insights into the dynamic regulatory landscape of cellular differentiation. Looking forward, this system additionally presents the opportunity to further explore these diverse contexts under a wide range of chemical and genomic perturbations, to better understand the role of gene-environment and gene-gene interactions. This expanded view of gene regulation offers a foundation to more deeply understand the genetics of complex traits, with the ultimate goal of accelerating the discovery of fundamental disease mechanisms and identifying potential therapeutic targets.

## Acknowledgments

We would like to thank Natalia Gonzales for comments on the manuscript, and Peter Carbonetto, Benjamin Strober, and Rebecca Keener for discussions. This work was supported by resources provided by the University of Chicago’s Research Computing Center and the Advanced Research Computing at Johns Hopkins University (ARCH) core facility.

## Author contributions

Conceptualization: AB, YG

Methodology: JMP, KR, KB, AB, YG

Formal Analysis: JMP, RJ, ML, KT, KB

Investigation: KR, KB

Funding acquisition: AB, YG

Supervision: AB, YG

Writing – original draft: JMP, AB, YG

Writing – review & editing: JMP, KR, RJ, ML, KB, KT, AB, YG

### Data and code availability

Raw sequencing data is available in the SRA under BioProject PRJNA1090081. Processed expression data has been submitted to GEO. Genotype data for the Yoruba of Ibadan, Nigeria population is available through the International Genome Sample Resource at https://www.internationalgenome.org/data-portal/population/YRI. Code needed to reproduce the analyses described in this paper is available in a snakemake workflow at https://github.com/jmp448/hdcQTL.

## Supplementary Materials

Document S1. Figs. S1 to S6

Document S2. Supplementary Tables S1 to S13

## Methods

### iPSC Maintenance

We maintained feeder-free iPSC cultures on Matrigel Growth Factor Reduced Matrix (CB-40230, Thermo Fisher Scientific) with StemFlex Medium (A3349401, Thermo Fisher Scientific) and Penicillin/Streptomycin (30,002 Cl, Corning). We grew cells in an incubator at 37 °C, 5% CO2, and atmospheric O2. Every 3–5 days thereafter, we passaged cells to a new dish using a dissociation reagent (0.5 mM EDTA, 300 mM NaCl in PBS) and seeded cells with ROCK inhibitor Y-27632 (ab120129, Abcam).

### Unguided HDC Differentiation

HDCs were formed using a modified version of the STEMCELL Agrewell400 protocol which was previously described in Rhodes et al. 2022. We coated wells of an Aggrewell 400 24-well plate (34414, STEMCELL) with anti-adherence rinsing solution (07010, STEMCELL). iPSCs were seeded into the Aggrewell 400 24-well plate at a density of 1,000 cells per microwell in Aggrewell EB Formation Medium (05893, STEMCELL) with ROCK inhibitor Y-27632 and Penicillin/Streptomycin. After 24 hours, we replaced half of the spent media with fresh Aggrewell EB Formation Medium without ROCK inhibitor. 48 hours after seeding the Aggrewell plate, we harvested EBs and moved them to an ultra-low attachment 6-well plate (CLS3471-24EA, Sigma) in E6 media (A1516401, ThermoFisher Scientific) with Penicillin/Streptomycin. We maintained HDCs in culture for an additional 19 days, replacing media with fresh E6 media every 48 hours.

HDCs were dissociated for collection 21 days after formation. HDCs were dissociated by washing them with phosphate-buffered saline (Corning 21-040-CV), treating them with AccuMax (STEMCELL 7921) and incubating them at 37C for 15-40 minutes total. After the first 10 minutes in Accumax, we pipetted HDCs up and down with a wide-bore p1000 pipette tip for 30 seconds. Subsequently, we repeated pipetting with a standard p1000 pipette tip for 30 seconds every 5 minutes until HDCs were completely dissociated. We then quenched the dissociation by adding E6 media to cells and strained them through a 40 um strainer (Fisherbrand 22-363-547). We resuspended cells in PBS with 0.04% bovine serum albumin and counted them. Early collection batches were counted using a TC20 Automated Cell Counter (450102, BioRad), and later batches were counted using a Countess II (AMQAF1000, invitrogen). We then mixed lines together in equal proportions prior to collection.

We differentiated two replicates of 51 iPSC lines across 17 batches consisting of 4 to 8 lines per batch. Each batch contained at least one line of each sex. In each batch, lines were formed and maintained in parallel. Each replicate represents a separate instance of HDC differentiation and will capture technical effects introduced during formation, maintenance, dissociation, and collection. We refer to these batches as “collection batches”.

In addition to these 51 iPSC lines which were differentiated and collected for this study, analyzed 3 batches previously collected in Rhodes et al. 2022, which included 2 additional lines, bringing the total number of cell lines in this study to 53.

### Single-cell sequencing

We generated scRNA-seq libraries using the 10X Genomics 3’ scRNA-seq v3.1 kit. Using the evenly pooled mix of lines from each collection batch, we loaded a 10x chip targeting 10,000 cells per lane of the 10x chip and loaded the same pool of cells across multiple lanes to recover 10,000 per individual. After cDNA amplification and cleanup, samples from each collection batch were stored at -20° C. Library preparation proceeded in larger batches composed of two or more collection batches (see Supplementary Table S1). For example, in library preparation batch 1, all samples from collection batches 1 and 3 were performed in parallel. cDNA libraries for biological replicates were always processed in different library preparation batches. Libraries were sequenced on the NovaSeq in the University of Chicago Functional Genomics Core. We pooled samples for sequencing in 7 batches. These batches are distinct from library preparation batch and are composed of two or more samples from the same collection batch. Some samples from a single collection day were split across different “sequencing batches” (Supplementary Table S1). We targeted a final sequencing depth of 100,000 reads per cell.

### Alignment, demultiplexing, and preliminary quality control

We used Cellranger to align samples to the human genome (GRCh38) and to aggregate samples from all collections from this study as well as additional libraries collected previously that included 2 additional Yoruba individuals (NA19160, NA18511) and one individual represented in both collections (NA18858) ^24,25,53^. We used Vireo to demultiplex samples and assign droplets to individuals. We used previously collected and imputed genotypes for the included Yoruba individuals from the HapMap and 1000 Genomes Project ^54–56^. We then filtered cells to remove droplets that Vireo identified as doublets and droplets that could not be confidently assigned to an individual. For the samples collected prior to this study, we kept only droplets representing Yoruba individuals (these samples originally included chimpanzee cells and cells from non-YRI humans). We further filtered cells to keep only those with less than 15% mitochondrial reads and with at least 2500 genes expressed. We also removed cells with very high total counts, keeping only those with less than 150,000 total counts. Finally, we filtered to genes expressed in at least 10 cells. This left a total of 909,536 cells and 35,324 genes for downstream analysis.

### Fetal cell atlas classifier

The fetal cell atlas contains a total of 77 cell-type labels. To obtain marker gene sets for these 77 cell-types, we subsampled overabundant cell-types to a maximum of 5,000 cells per cell-type (regardless of the tissue of origin), and filtered genes to protein-coding genes. To assess classifier performance, we treated cell-type labels assigned by the original fetal cell atlas paper as ground truth. We split the uniformly sampled data into training and testing subsets with equal numbers of cells. We preprocessed the training data using *scater* ^57^: we first normalized cells by size factors, then log transformed the data, and selected highly variable features at an FDR threshold of 0.1. We then performed multiple correspondence analysis (MCA) to extract signature gene sets for each cell-type using *Cell-ID* ^58^. Annotation of the HDC data with these gene sets suggested that 64 of 77 cell-types were represented with at least 5 cells present from at least 25 donors. However, this classifier displayed poor accuracy on held-out test data (Fig. S1). Examination of the reference gene sets suggested that high similarity among related cell-types (e.g., neuronal subtypes) likely compromised the classifier’s performance.

In order to obtain a more limited set of maximally interpretable cell-type labels, we first removed 12 cell-types which were poorly characterized in the reference dataset, or attributed to potential contamination: AFP_ALB positive cells, CCL19_CCL21 positive cells, CLC_IL5RA positive cells, CSH1_CSH2 positive cells, ELF3_AGBL2 positive cells, MUC13_DMBT1 positive cells, PDE11A_FAM19A2 positive cells, PDE1C_ACSM3 positive cells, SATB2_LRRC7 positive cells, SKOR2_NPSR1 positive cells, SLC24A4_PEX5L positive cells, and SLC26A4_PAEP positive cells. We additionally removed placental cells from the reference which are unlikely to arise in the *in vitro* HDC system, as well as rare cell-types represented by fewer than 500 cells for which it may be difficult to define a meaningful expression signature. We combined limbic system neurons, inhibitory interneurons, inhibitory neurons, excitatory neurons, unipolar brush cells, granule neurons, and Purkinje neurons under the label of central nervous system (CNS) neurons; microglia, astrocytes, and oligodendrocytes under CNS glia; visceral neurons and enteric nervous system (ENS) neurons under peripheral nervous system (PNS) neurons; ENS glia, satellite cells, and Schwann cells under PNS glia; retinal progenitors and Muller glia, amacrine cells, bipolar cells, ganglion cells, retinal pigment cells, horizontal cells, and photoreceptor cells under retinal cells; and chromaffin cells, Islet endocrine cells, neuroendocrine cells, and sympathoblasts under neuroendocrine cells. Immune cell-types were also quite granularly defined in this reference, posing a challenge to a global cell-type classifier. We therefore removed myeloid cells and hematopoietic stem cells from the reference, which are common ancestors of more specific cell-types present in the reference, and merged thymocytes with lymphoid cells due to their shared lymphoid progenitor. Altogether, this left us with a refined set of 33 cell-types. After once again subsampling overrepresented cell-type labels to at most 5000 cells, and re-defining our set of highly variable genes using the same criteria as above, classification accuracy increased to nearly 90% (Fig. S1).

Our final set of cell-type signatures was obtained by applying the sampling, normalization, and embedding procedures described above to the merged training and test datasets. The gene set signatures used for this final classifier are available in Supplementary Table S2.

### Cell-type annotation

To classify HDC cells using these gene sets, we first normalized our HDC cells as described in the previous section, subsetting to the same set of highly variable genes used for the fetal cell atlas embedding. We applied MCA, and used *Cell-ID* to generate cell-type labels.

### UMAP visualization

To generate the UMAP embedding, we first identified 5000 highly variable genes using proportional fitting ^59^ and scanpy’s default method for extracting highly variable genes proposed in ^60^. We subset to these 5,000 genes to generate a 50-dimensional embedding of the data using a variational autoencoder applied to raw UMI counts with *scVI* ^61^. To generate a 2-dimensional embedding of the data we computed a neighborhood graph based on this encoding and applied UMAP using *scanpy*’s default settings ^62^. This UMAP embedding was used purely for the convenience of visualizing the full dataset in two dimensions, and did not influence cell-type annotation, trajectory inference, topic modeling, or any QTL calling.

### Hierarchical clustering of cell-types

To group cell-types according to expression similarity, we aggregated all cells with the same cell-type label into a pseudobulk sample by taking the sum of all UMI counts. We then filtered to the intersection of protein-coding genes and the fetal cell atlas highly variable gene set (see *Classifier Development* section), and applied TMM and log CPM normalization using the *edgeR* package ^63^. We applied hierarchical clustering to these normalized pseudobulk samples for each cell-type using Ward’s method ^64^.

### eQTL calling in each cell-type

To perform eQTL calling, we aggregated cells from the same donor and cell-type by taking the sum of all UMI counts per gene. We removed pseudobulk samples with fewer than 5 cells. Within each cell-type, we filtered to genes with nonzero variance, and with a median expression level of at least 10 UMI counts per sample. We applied log CPM normalization to each sample, using TMM normalization factors computed separately in each cell-type with the *edgeR* package ^63,65^. We then applied an inverse normal transformation to each gene. We computed expression principal components to include as latent covariates for eQTL calling, as well as for quality control: principal component biplots were manually inspected to identify and remove outlier samples which may disrupt eQTL calling. Results of these filtering criteria are available in Supplementary Table S3.

We tested all variants within 50kb of the corresponding gene’s TSS, that had minor allele frequency of at least 0.1 among specifically the samples included in each cell-type specific analysis (note that this set of donors varies between cell-types as demonstrated in Figure 1D). Genotypes were normalized by a factor of 1/[p(1-p)] (where p is the minor allele frequency) to maximize comparability of effect size estimates across cell-types. (This does not influence significance tests for the cell-type-by-cell-type analysis, but becomes relevant for the mash analysis described below). QTL calling was conducted with TensorQTL, using beta-approximated *P*-values based on permutations to control for multiple testing burden at each gene^66^.

To generate a list of all significant variant-gene pairs, we followed the procedure described by the GTEx Consortium ^4^: a genome-wide *P*-value threshold was defined as the beta-approximated *P*-value of the gene closest to the global q-value cutoff of 0.05. This genome-wide threshold was used to define a nominal threshold for each gene based on the per-gene beta distribution estimated with TensorQTL, and all variants with a nominal *P*-value below this gene-level threshold were considered significant (Supplementary Table S5).

### GTEx overlap

We used bedtools ^67^ to identify variant-gene pairs intersecting the full set of significant eQTL variant-gene pairs in each GTEx tissue, using data from GTEx v8.

### Multivariate adaptive shrinkage (MASH) analysis

To perform the mash analysis, we first estimated residual correlation structure (V) with mash’s expectation maximization procedure, subsetting to a random subset of 25,000 gene-variant pairs that were tested in all cell-types for efficiency ^68^. Next, to estimate data-driven covariance matrices (U), we subset first to gene-variant pairs which were tested in all cell-types, then to the strongest (largest absolute z-score) effect per gene, and finally to the top 2,000 strongest effects across all genes. We decomposed this set of 2,000 eQTL effects across cell-types into non-negative latent factors through Empirical Bayes Matrix Factorization using flashier ^69^ as implemented in the mash package. We removed singleton components which only assigned weight to a single cell-type. We included the four rank-1 covariance matrices generated from these latent factors, as well as their normalized sum, to fit the mash model. We additionally included a singleton component for each cell-type as well as a component corresponding to shared effects across all cell-types. We fit the mash model on a random subset of 50,000 gene-variant pairs that were tested in at least 10 contexts. After model fitting, we performed inference across all variant-gene pairs tested in at least 5 contexts.

### Trajectory isolation

For each trajectory, we curated marker gene lists from multiple published directed differentiation protocols defining marker genes for each stage of differentiation ^17,70–106^. We used scDRS to test for enrichment of the marker gene set compared to 50 background gene sets matched for mean and variance of library-size normalized, log-transformed single-cell expression ^36^, filtering to cells with FDR <= 0.1 for inclusion in the isolated trajectory. Marker gene sets are available in Supplementary Table S7-S9.

### Pseudotime inference and marker gene visualization

We applied principal component analysis to library-size, log normalized single-cell gene expression data from each trajectory in isolation, and used this first principal component as a measure of pseudotime. To visualize trends in marker gene expression with respect to pseudotime, we adopted a sliding window approach used by Cuomo et al (taking the average expression of 10% of cells in each window, sliding along pseudotime by 2.5% of cells) ^15^, followed by min-max normalization for each gene.

### Pseudobulk aggregation

To mitigate the noise of single-cell data and leverage efficient and established tools for interaction eQTL calling, we aggregated cells into pseudobulk samples. We grouped cells into 15 pseudotime bins of equal range. In each trajectory, cells from bins 1 and 2 were combined as were cells from bins 14 and 15 to account for fewer cells and donors being represented at the extremes of the pseudotime distribution. Each pseudobulk sample contains the sum of UMI counts from all cells from a single donor in a single pseudotime bin.

### Pseudobulk sample, gene, and variant filtering for dynamic eQTL calling

In each trajectory, we dropped pseudobulk samples that consisted of less than 5 cells, omitted any donors represented in 10 or fewer pseudotime bins, and filtered to genes with non-zero pseudobulk expression in at least 10 samples and non-zero variance in expression across samples. This resulted in a total of 428 pseudobulk samples from a total of 35 cell lines for the cardiomyocyte trajectory, 383 samples from 31 cell lines in the hepatocyte trajectory, and 549 samples from 44 cell lines in the neuronal trajectory. We normalized pseudobulk expression and filtered tests as described for the cell-type analysis.

### Covariates for dynamic eQTL calling

As in previous work ^17,21^, we used cell line PCA to infer latent covariates introducing broad differences in expression between cell lines over the course of a differentiation trajectory. We used the NIPALS algorithm ^107^ to perform cell line PCA with missing values, as not all donors contain 5 cells in each pseudotime bin. We included sex and the first 10 cell line PCs as covariates, as well as their interaction with pseudotime.

### Dynamic eQTL calling

We use TensorQTL ^66^ to conduct dynamic eQTL calling, using the following model:

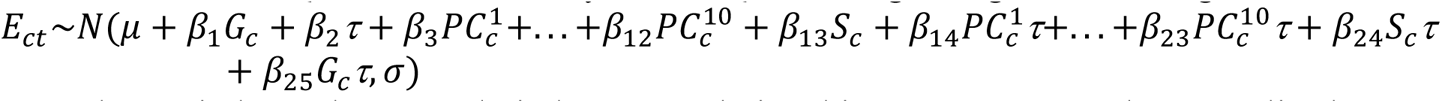

Where *c* indexes donors and *t* indexes pseudotime bins. *E_ct_* represents the normalized expression of the sample, *G* represents genotype, *PC^k^* represents the k^th^ cell line PC, *S_c_* represents the sex of donor *c*, and *τ* represents median pseudotime value of all cells in the sample.

We perform inference on the interaction effect between genotype and pseudotime (β_25_). We used eigenMT to adjust for multiple QTL tests conducted per gene ^108^. We then apply the Benjamini-Hochberg procedure to the smallest adjusted *P*-value per gene to control the genome-wide false discovery rate at a level of 0.1.

We generated a list of all significant dynamic eQTLs analogously to the procedure described for cell-type eQTLs: a genome-wide adjusted *P*-value threshold was defined as the EigenMT-adjusted *P*-value of the gene closest to the global FDR threshold, and all variants with an adjusted *P*-value below this threshold were included in the final set of significant dynamic eQTLs (Supplementary Table S10).

### Replication of cardiomyocyte dynamic eQTLs

To test for replication of these eQTLs in a separate dataset, we reprocessed the single-cell expression data collected in Elorbany et al as described above, and used the *qvalue* package to estimate the π_1_ replication rate for the strongest cardiomyocyte dynamic eQTL per gene detected in our study ^33^.

### Classification of dynamic eQTLs

We classified significant dynamic eQTLs as early, late, or switch categories as in previous work ^17,21^. An eQTL is classified as early if the effect size decreased over time, late if it increased over time, and switch if the sign of the effect changed over time. We used the fitted linear model generated by TensorQTL to estimate predicted effect sizes at the endpoints of the trajectory’s pseudotime range, and compare these predictions to classify each eQTL. If the effect size remains the same, the variant is classified as early if the magnitude decreases or late if the magnitude increases. If the effect size flips and the predicted effect at both ends has a magnitude exceeding a threshold of 1, it is called a switch effect.

### Pseudocell aggregation for topic modeling

To improve computational efficiency and once again mitigate the noise of single-cell data, we aggregated cells into pseudocells before applying topic modeling in an approach similar to that used by Strober et al ^109^. To avoid over-representation from donors with outlier cell counts, for donors with cell counts over 1.5 standard deviations above the median we randomly subsampled without replacement to the median number (16,839) of cells. We also removed cells collected in a previous study due to differences in cell depth that disrupted topic modeling, leaving 51 of 53 donors. For each combination of donor and collection batch, we generated a separate neighborhood graph based on the *scVI* embedding space (see *UMAP visualization*) and applied Leiden clustering, as implemented in the *scanpy* package, at resolution 15. We summed raw UMI counts for all cells within a cluster to generate pseudocell expression. This left 17,913 pseudocells after aggregation, with a median of 37 cells per pseudocells and a median of 342 pseudocells per donor.

### Topic modeling

To fit the topic model, we first filtered out genes expressed in less than 10 pseudocells. We then applied Poisson negative matrix factorization to the filtered pseudocell expression data: we used k=10 topics, and fit the model first using 400 expectation maximization (EM) updates followed by 200 stochastic coordinate descent (SCD) updates ^42^. We then use the parameter estimates from this Poisson non-negative matrix factorization to recover parameter estimates for the multinomial topic model using *FastTopics*’s poisson2multinom command.

### Topic interpretation

We conducted grade of membership differential expression (DE) analysis using *FastTopics*’s de_analysis function. Genes with posterior log fold change values over 2 were considered for gene set enrichment using the Gene Ontology ^110^. We performed a hypergeometric test for topic driver gene enrichment in all GO biological processes, and used the Benjamini-Hochberg procedure to control the false discovery rate at a level of 0.01. We focused on overrepresentation of topic DE genes in a gene set for the purposes of interpretation (OR >= 1, Supplementary Table S11). Topics 3 and 9 displayed no enrichment for any GO biological processes nor clear enrichment in any of the previously characterized cell-types, leading us to omit these two topics from downstream QTL calling analysis to instead focus on the 8 more clearly interpretable latent topics.

### Topic eQTL calling

CellRegMap takes as inputs a kinship matrix to account for genetic similarity between cells, an environmental covariance matrix to account for similarity due to cellular context, and covariates. We used *Plink* ^111^ to construct the kinship matrix from all 53 donors. To construct the environmental covariance matrix, we applied an inverse normal transformation to each column (topic) of the topic loadings matrix (C, pseudocells x topics), then did the same transformation for each row (pseudocell), before generating the environmental covariance matrix (CC^T^). We included sex and collection date as covariates. After QTL calling, we applied Bonferroni correction and generated a list of significant topic eQTLs as in both previous QTL analyses: a genome-wide adjusted *P*-value threshold was defined as the Bonferroni-adjusted *P*-value of the gene closest to a global q-value threshold of 0.1, and all variants with an adjusted *P*-value below this threshold were included in the final list (Supplementary Table S12).

### Schizophrenia GWAS overlap

To identify schizophrenia risk variants with novel regulatory interactions identified in HDCs, we filtered the aggregate list of significant variant-gene pairs from all cell-types to eQTLs that did not overlap an eQTL in any tissue according to the GTEx v8 Catalog. We then intersected these variants with genome-wide associated risk variants (P <= 5e-8) from Pardinas et al ^112^. For this set of genome-wide significant HDC eQTLs without GTEx overlap, we compiled a list of all variants within 1Mb of the SNP that was in LD at R2 >= 0.5 (using the 53-donor cohort in this study to compute LD). We then additionally checked these tag variants for overlap with any GTEx eQTLs in any tissue, and removed any such GTEx-overlapping tag variants to obtain our final set of schizophrenia-associated HDC eQTLs with no GTEx overlap.

### Phenome-wide effects of interaction eQTLs

To identify phenotypic effects of interaction eQTLs, we used an approach similar to that described for schizophrenia overlap. Instead of cell-type eQTLs, we used the aggregate set of significant variant-gene pairs from all three trajectory dynamic eQTL analyses, as well as the significant variant-gene pairs from the CellRegMap analysis. We subset to interaction eQTLs that did not overlap a GTEx eQTL in any tissue, and queried the OpenTargets database for any GWAS studies in which these interaction eQTLs displayed genome-wide significant effects. We similarly removed any tagged variants with known regulatory effects in the GTEx Catalog to obtain a final set of GWAS-overlapping HDC eQTLs with no GTEx overlap.

