## Supplementary Figures for "Cell-type and dynamic state govern genetic regulation of gene expression in heterogeneous differentiating cultures"

**The PDF file includes:**

Figs. S1 to S6

**Other Supplementary Materials for this manuscript include the following:**

Document S2. Supplementary Tables S1 to S13

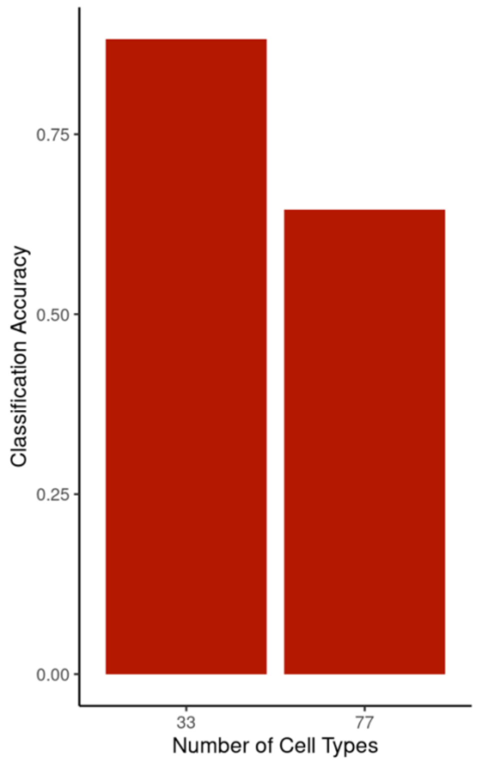


**Fig. S1:** Using original fetal cell atlas labels as ground truth, comparison of classification accuracy based on all 77 cell-types or a refined set of 33 cell-types.

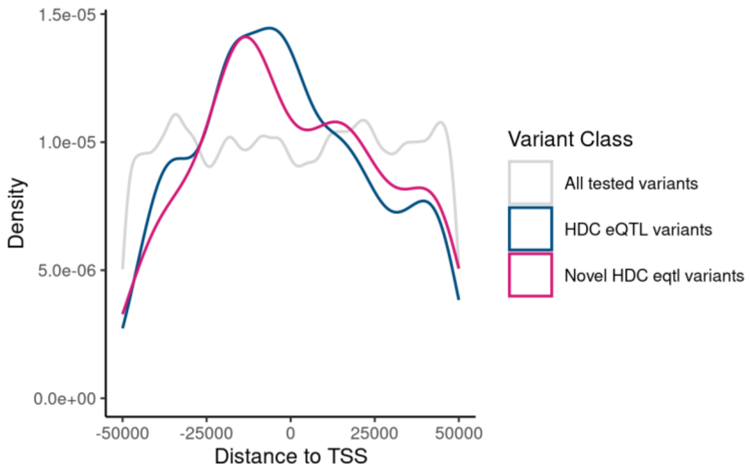


**Fig. S2:** Genomic distribution of all variants tested for eQTLs of tissue development genes (all tested variants, gray), the subset of these which had significant eQTL variants (HDC eQTL variants, dark blue), and the subset of these eQTL variants which were not detected anywhere in GTEx (novel eQTL variants, pink).

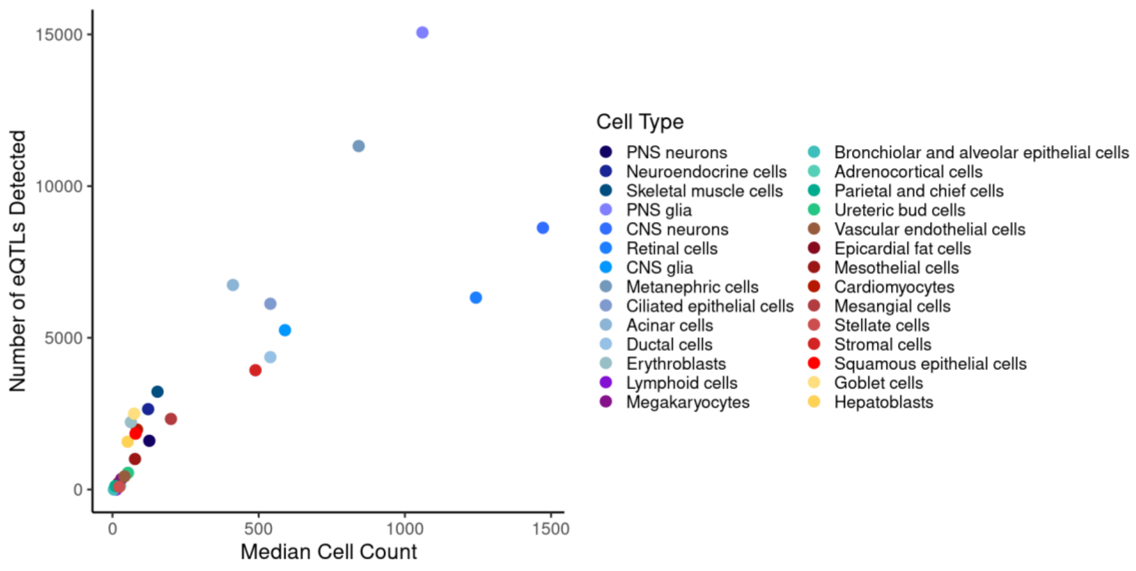


**Fig. S3:** Number of eQTLs detected per cell-type, compared to the median cell count across all samples.

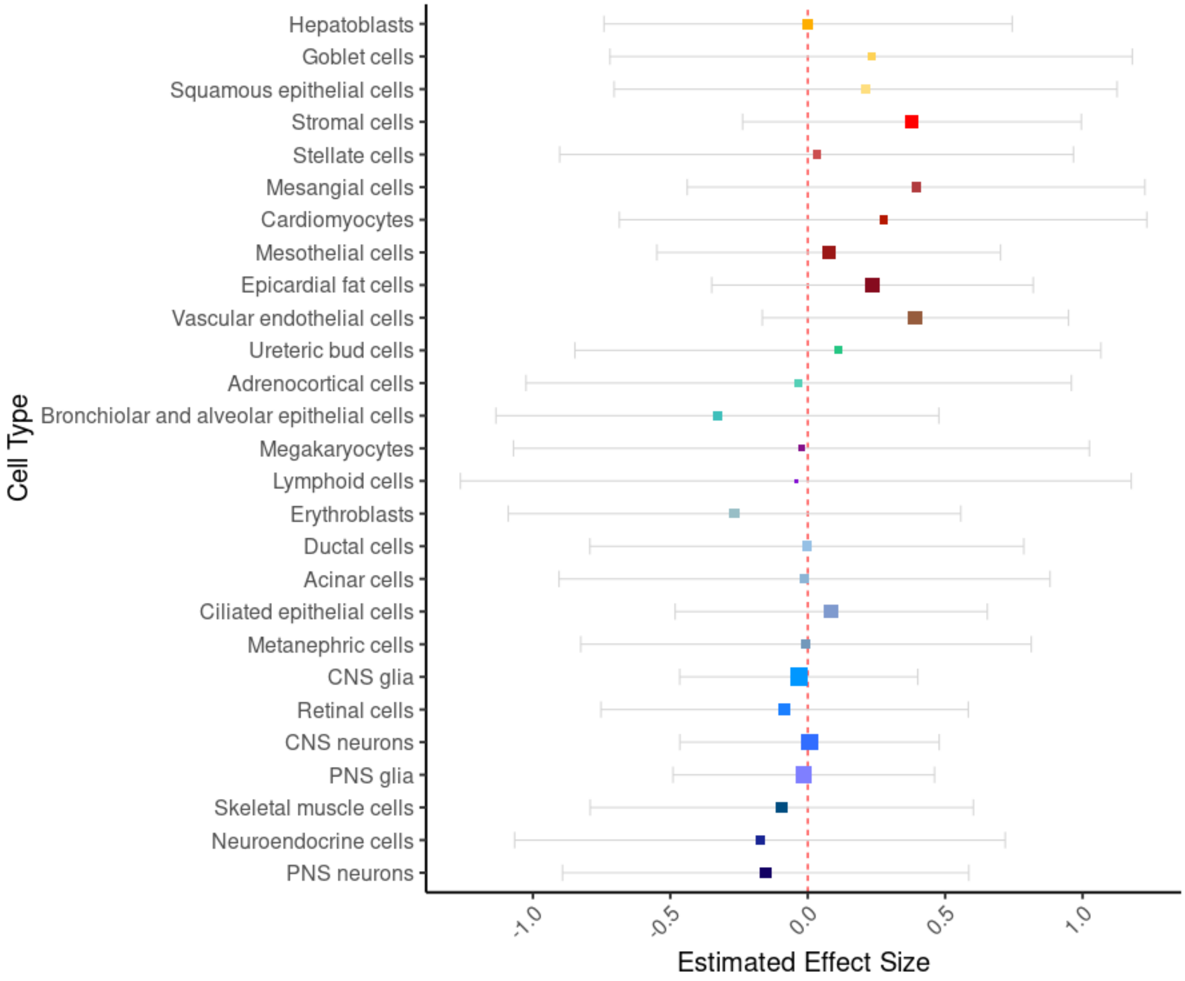


**Fig. S4:** eQTL effect estimates for the gene *SH3PXD2B* at rs10042482, based on eQTL calling in each cell type in isolation (no meta-analysis performed). Boxes are centered at the estimated effect size in the given cell type, error bars show +/- standard deviation, box size indicates precision (1/squared standard deviation).

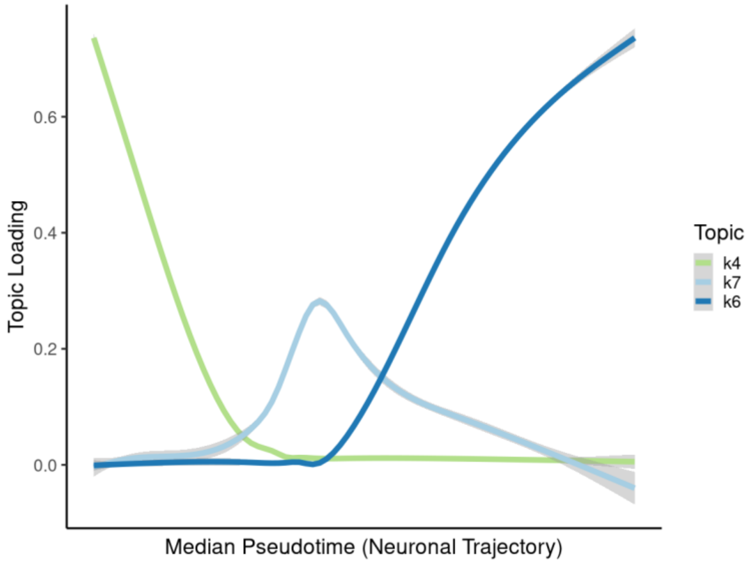


**Fig. S5:** Comparing pseudocell topic loadings to median pseudotime values along the neuronal trajectory highlight topics corresponding to early, intermediate, and late stages of neuronal differentiation.

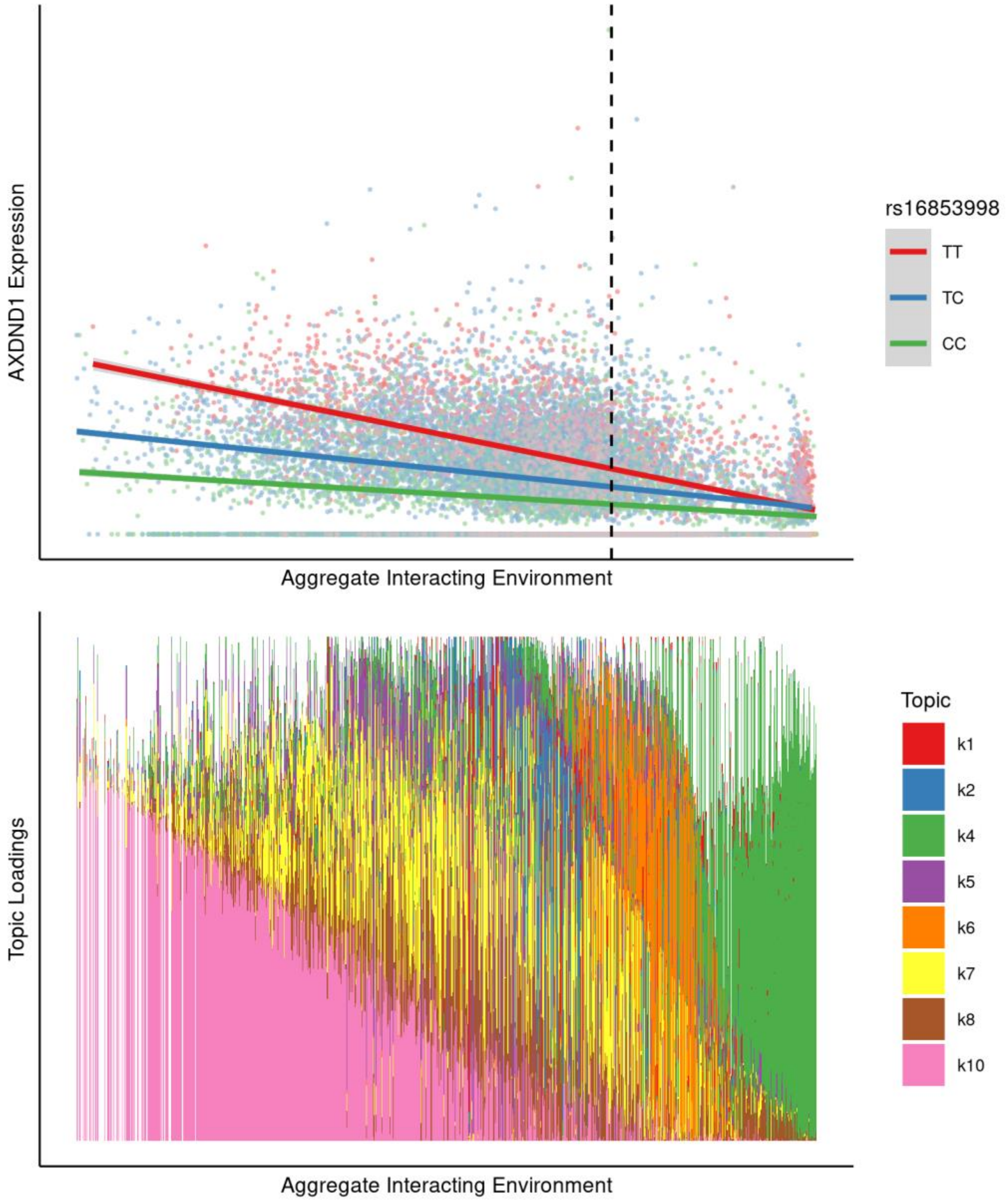


**Fig. S6:** Example topic eQTL for the gene *AXDND1* with maximal effect in cells highly loaded for topic 10 (pink), a topic associated with a ciliary gene program.
